# Rpl12 paralog dependent TOR-signaling controls the expression of ribosome preservation factor Stm1

**DOI:** 10.64898/2026.03.25.714144

**Authors:** Sourav Sharma, Pratyush Kumar Datta, Suresh Singh Yadav, Imran Pancha, Rohini R Nair

## Abstract

Ribosomes are increasingly recognized as heterogeneous regulators of gene expression, yet how ribosomal protein paralogs interface with nutrient signaling remains poorly understood. In *Saccharomyces cerevisiae*, ribosomal protein gene expression is governed by duplicated gene pairs, many of which exhibit functional divergence despite high sequence identity. A central regulator of ribosome biogenesis and translational control is the Target of Rapamycin (TOR) pathway, which integrates nutrient signals to modulate growth, stress adaptation, and lifespan. Target of Rapamycin Complex 1 (TORC1) influences ribosome activity by phosphorylating ribosomal protein S6 (Rps6), a modification that links nutrient availability to translational output.

Here, we investigated the functional divergence of the Rpl12 ribosomal stalk protein paralogs *RPL12a* and *RPL12b* and found that *rpl12bΔ* produces phenotypes consistent with reduced TOR activity, including decreased Rps6 phosphorylation, G2/M cell-cycle accumulation, and significant extension of chronological lifespan as compared to *rpl12aΔ* and wildtype strain. Multi-omics analyses further indicate translational and metabolic reprogramming consistent with activation of a stress-adaptive program associated with Gcn4. Importantly, loss of *RPL12b* also reduces levels of the ribosome preservation factor Stm1, a TORC1-regulated protein required for stabilization of 80S ribosomes under stress. This finding links ribosomal stalk composition to ribosome stability and nutrient-responsive signaling. Together, our results demonstrate that ribosomal paralog specialization provides an additional regulatory layer connecting translation, TOR signaling, and cellular longevity.

## INTRODUCTION

Ribosomes previously thought of being a passive, homogeneous, indiscriminate molecular machine have recently been transitioned into highly heterogeneous and macromolecular complexes with specialized functions (Genuth and Barna 2018; Aspden et al. 2025). Despite this conservation, their structural and functional characteristics vary significantly between prokaryotes and eukaryotes, reflecting their adaptation to the complexity of the cellular environment (Melnikov et al. 2012). Initially termed “microsomes” in 1955, ribosomes were once considered passive and uniform translation machines (Palade 1955), however with the consistent structural and functional variation lead to the emergence of ribosomal heterogeneity and specialized cellular functions (Crick 1958; Genuth and Barna 2018). Ribosomal heterogeneity may arise due to stoichiometric difference of ribosomal proteins on the ribosome, variations in Ribosomal Proteins (RPs) composition, presence of paralogs of Ribosomal Proteins (RPs), post-translational modification (PTMs) of RPs, variation in translation initiation factors, rRNAs or tRNAs and may be contributed by other aspects which involves differential interaction with ribosomal associated proteins or their subcellular localization (Sauert et al. 2015; Genuth and Barna 2018; Li and Wang 2020).

The *S. cerevisiae,* whole-genome duplication preserved 59 ribosomal protein paralog pairs, likely due to substantial demand for ribosomal protein synthesis. However, studies show that these paralogs are not fully redundant, instead contributing distinct regulatory roles in translation and metabolism (McIntosh and Warner 2007; Komili et al. 2007). One example is Rpl12, the eukaryotic homolog of bacterial uL11, which stabilizes the ribosomal GTPase domain and supports elongation factor interactions (Briones et al. 1998). In bacteria, *uL11* interacts with proteins *L10* and *L7/L12* to form the ribosomal stalk, a pentameric complex required for GTP hydrolysis during elongation (Stark et al. 1997). Eukaryotic counterparts include P0 (L10 homolog), P1 (L7 homolog), and P2 (L12 homolog) which assemble to form a pentameric complex P0-(P1)₂-(P2)₂, that associates with Rpl12 *(uL11 homolog)* to finally form the ribosomal stalk (Uchiumi et al. 1987; Mager et al. 1997). In *yeast*, Rpl12 is encoded by two paralogs, *RPL12a* and *RPL12b*, both required for optimal growth (Briones et al. 1998; García-Marcos et al. 2007), yet their specific contributions to translational control and stress adaptation remain unresolved. Apart from the structural role, Rpl12 also has a regulatory role. Recent phosphoproteomics studies have shown that Rpl12 is phosphorylated at serine 38 (S38) during mitosis by the action of CDK1-cyclin B complexes. This phosphorylation event, although not affecting protein synthesis globally, increases the translation of mitosis-related mRNAs (Imami et al. 2018).

Moreover, a recent study describes that another function of Rpl12 is that it serves as a conserved receptor for ribophagy. Under starvation conditions, it is phosphorylated at Ser79 and Ser101 by Atg1 kinase, thereby increasing its interaction with the scaffold protein Atg11 and enhancing the recruitment of the ribosomes to the autophagosomes. Loss of Rpl12-mediated ribophagy reduces survival under nutrient stress and pathogen infection in both yeast and C. elegans. These characteristics highlight the physiological significance of ribophagy in stress resistance, immunity, development, and longevity (Chen et al. 2025). These findings suggest that Rpl12 is not only structural but also participates in ribosome turnover and stress resistance. How its paralogs integrate with nutrient-sensing pathways, particularly the Target of Rapamycin (TOR) pathway, is a key open question.

Ribosome specialization interfaces directly with nutrient-sensing networks, especially the Target of Rapamycin (TOR) signaling ,TOR is a conserved Ser/Thr kinase that forms two complexes in yeast: TORC1 (consists of either Tor1 or Tor2 and other proteins) and TORC2 (consists of only Tor2 and other proteins) (Ikai et al. 2011). Being a hub protein, TORC1 is involved in cell growth, ribosome biogenesis, translation initiation, metabolism, stress response, aging, and autophagy whereas TORC2 is involved in actin organization, sphingolipid biogenesis, and endocytosis (Loewith et al. 2002; González and Hall 2017). Rapamycin is a potent inhibitor of the Target of Rapamycin Complex 1 (TORC1) used to uncover the role of TORC1 as a master regulator for cellular growth. Under nutrient-rich conditions, TORC1 promotes ribosome biogenesis and protein synthesis, while its inhibition through rapamycin suppresses these processes and activates stress-responsive programs that conserve energy and reduced phosphorylation of ribosomal protein S6 (Rps6) downstream TOR signaling pathway directly linking nutrient availability directly to ribosome function and enhanced survival (T. Powers and Walter 1999; Jacinto and Hall 2003; R. W. Powers et al. 2006; Rohde et al. 2008).

In yeast, chronological lifespan (CLS) is strongly influenced by nutrient-sensing pathways. The TOR-Sch9 pathway is a major pro-aging pathway, and its inhibition also extends lifespan by activating transcription factors Msn2/4, which control oxidative stress resistance, and Gis1, which is a regulator of metabolic adaptation during the post-diauxic shift (Wei et al. 2008; Deprez et al. 2018). In parallel, the transcription factor Gcn4 is activated during amino acid starvation and triggers amino acid biosynthesis, autophagy, and stress resistance pathways that contribute to longevity. (Steffen et al. 2008a; Mittal et al. 2017b; Gulias et al. 2023a). Thus, inhibition of the TOR-Sch9 pathway and activation of the Gcn4 pathway converge to induce metabolic adaptations that provide stress resistance, nutrient homeostasis, and longevity.

Additionally, ribosome associated factors contribute to TOR regulated adaptation. Stm1, a ribosome preservation factor in *S. cerevisiae*, binds inactive 80S ribosomes under nutrient limitation or TORC1 inhibition, stabilizing them in a dormant state and protecting against degradation. Upon nutrient repletion, active TORC1 phosphorylates Stm1, releasing ribosomes for translation re-initiation. This regulation positions Stm1 as a sensitive readout of ribosome-nutrient pathway crosstalk, highlighting how TORC1 controls not only ribosomal protein phosphorylation (e.g., Rps6) but also ribosome-associated factors (Yerlikaya et al. 2016; Shetty et al. 2023).

To determine whether Rpl12 paralogs generate specialized ribosomes that influence cellular translation and nutrient signaling, we characterized the *rpl12aΔ* and *rpl12bΔ* mutants with respect to growth, TOR signaling, translational and metabolic regulation. We observed that although both mutants showed only modest growth defects under nutrient-rich conditions, deletion of Rpl12b produced pronounced phenotypes under and similar to TOR inhibition. We observed *rpl12bΔ* cells exhibited impaired growth in the presence of rapamycin, reduced phosphorylation of ribosomal protein Rps6, accumulation of cells in the G2/M phase and extended lifespan suggests Rpl12b is required to maintain proper ribosomal stoichiometry and growth regulation to maintain cellular homeostasis. Moreover, translatome mapping using a nascent chain sequencing approach (puromycin-associated nascent chain proteomics; PUNCH-P;(Singh et al. 2024) and metabolic analysis showed that the differential regulation of genes involved in amino acid biosynthesis and metabolic pathways in *rpl12bΔ* relative to *rpl12aΔ*. Several genes associated with the amino acid starvation response and known targets of the transcription factor Gcn4 were preferentially upregulated in *rpl12bΔ* cells.

Together, these findings suggest that Rpl12 paralogs differentially influence translational regulation and metabolic adaptation. Loss of *RPL12b* appears to promote a stress-adaptive translational program associated with reduced TOR activity and extended lifespan, supporting the concept that ribosomal protein paralogs can generate functionally specialized ribosomes that selectively shape cellular physiology

## RESULTS

### Rpl12 paralogs confer distinct growth responses under TORC1 inhibition

In the present study, we assessed the growth rate of Rpl12 paralogs, wild-type (WT), *rpl12aΔ* and *rpl12bΔ* mutant in defined medium (YPD) and we found Rpl12 paralog deletion *rpl12aΔ* (1.45 h, p < 0.0001) and *rpl12bΔ* (1.32 h, p < 0.01) has moderately reduced growth rate compared to the WT strain (1.2 h). (**Fig. 1A, B**). This reduction likely reflects the role of Rpl12 as a ribosomal stalk protein might be required for elongation factor interaction and ribosomal translocation, where deletion of either paralog impairs cellular growth.

**Figure 1.**
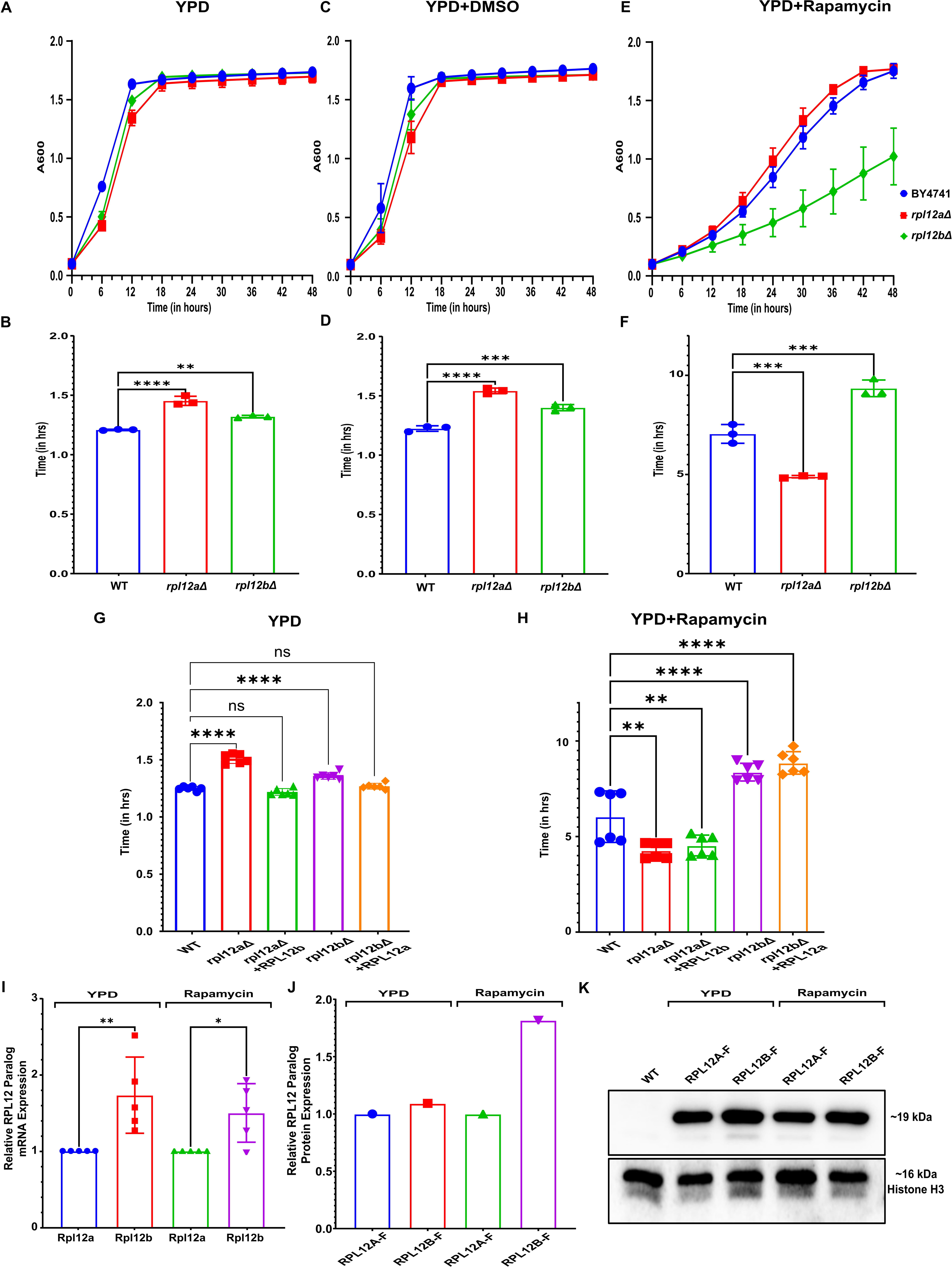
*RPL12* paralog mutants display show resistant and sensitive phenotype under rapamycin condition. **(A–F)** Growth kinetics of wild-type BY4741, *rpl12aΔ*, and *rpl12bΔ* strains in YPD medium alone (A–B), YPD + DMSO **(C–D)**, and YPD + rapamycin **(E–F)**. Growth curves **(A, C, E)** show absorbance at 600 nm over time, and corresponding bar graphs **(B, D, F)** depict doubling times were analysed using ordinary one-way ANOVA with indicated statistical significance according to adjusted P values. **(G–H)** Overexpression of paralogs does not compensate for the loss of the corresponding gene under rapamycin treatment. WT, RP deletion, and RP deletion strains overexpressing their corresponding a paralog from a single-copy plasmid were grown on YPD or YPD + rapamycin. Data in **(H)** were analyzed using ordinary one-way ANOVA with adjusted P values indicating statistical significance. **(I)** Relative mRNA expression levels of *RPL12a* and *RPL12b* under YPD and rapamycin-treated conditions, normalized to *HHF1* mRNA as a control. **(J–K)** Relative protein abundance and representative Western blot analysis of Rpl12a-Flag and Rpl12b-Flag under YPD and rapamycin-treated conditions, with Histone H3 used as a loading control. Error bars represent SEM.

However, under TOR inhibition, significant difference emerged: *rpl12aΔ* cells displayed a faster growth rate (4.8 h, p < 0.001) than wild type (6.93 h), whereas *rpl12bΔ* cells exhibited markedly slower growth (9.20 h, p < 0.001) (Fig1. E, F). These observations suggest paralog-specific functions in TOR-regulated growth, with *RPL12a* being more sensitive to rapamycin and Rpl12b supports growth under TOR inhibition. Simultaneously, the cells were also treated with DMSO (used to dissolve rapamycin) as control to check whether the growth rate is affected or not but we noticed it shows significantly similar growth comparable to YPD (**Fig. 1C, D**), confirming that the observed effects are rapamycin dependent.

To further understand the role of Rpl12 paralogs in TOR signaling, we exogenously overexpressed *RPL12b* and *RPL12a* in *rpl12aΔ* and *rpl12bΔ* mutants respectively under native promotor and terminator and checked the growth rate in YPD and the result indicates the restoration in growth similar to WT, as indicated by non-significant differences in generation time (∼1.22 h) **(Fig1 F)** similar to wild type supporting the paralog specific functional contribution. However, under TOR inhibition, overexpression failed to rescue the growth similar to WT i.e. showing growth similar to *rpl12aΔ* and *rpl12bΔ* mutants (**Fig.1 G)**. This confirms that the paralogs might confer non-redundance and become critical during TOR inhibition, revealing functional specialization of Rpl12 paralogs. This might contribute to ribosome specialization, where paralog composition may differentially modulate ribosome function during stress, potentially affecting selective mRNA translation rather than global protein synthesis.

This difference may arise from differential expressions of specific ribosomal protein paralogs or not by byproduct of alteration in gene expression, we measured mRNA level of *RPL12* paralogs in WT cells grown with and without rapamycin. We observed the significant increase in *RPL12b* (p < 0.01 and p < 0.05) mRNA expression as compared to *RPL12a* in YPD condition as well as under TOR inhibition, indication paralog specific transcription occurrence (**Fig.1 H)**. Simultaneously, we flag tagged *RPL12a* & *RPL12b* endogenously at C terminus in wild type and checked the protein level using anti-flag antibody. Immunoblot analysis further shows that the RPL12b expression is slightly higher as compared to RPL12a (**Fig.1 I, J)** i.e increased expression is maintained both at the transcription and translation level, confirming it as the dominant paralog.

Together, these findings suggest that preferential expression of RPL12b may contribute to maintaining ribosomal stoichiometry, particularly during TOR inhibition, thereby enabling adaptive regulation of ribosome composition and translational control.

### RPL12b deletion alters cell cycle progression and extends lifespan through impaired TOR signaling

To investigate whether impaired TOR signaling influences cell-cycle dynamics, we analyzed the cell-cycle distribution of WT, *rpl12aΔ*, and *rpl12bΔ* strains. Comparative cell cycle analysis reveals no such apparent difference exists between WT and *rpl12aΔ* paralog in yeast; however significant differences can be observed for *rpl12bΔ in* G2 phase (p < 0.05). WT and *rpl12aΔ* almost have a similar profile, with even distribution of cells in the G1, S and G2 phases suggests that cells are efficiently transiting through the cell cycle, with no obvious arrest or delay. This pattern indicates that the cells are healthy and cycling, with normal passage through the G1/S and G2/M stages. However, *rpl12bΔ* differs significantly from this pattern. It has significant accumulation in the G2/M phase (∼35%), with a non-significant reduction in the G1 phase (**Fig.2 A)**. This signifies a cell cycle arrest or delay at the G2/M transition, which could be caused by the G2/M checkpoint being activated. This suggests that Rpl12b plays an important role in the cell cycle progression, possibly by regulating the translation of cell cycle regulators.

**Figure 2.**
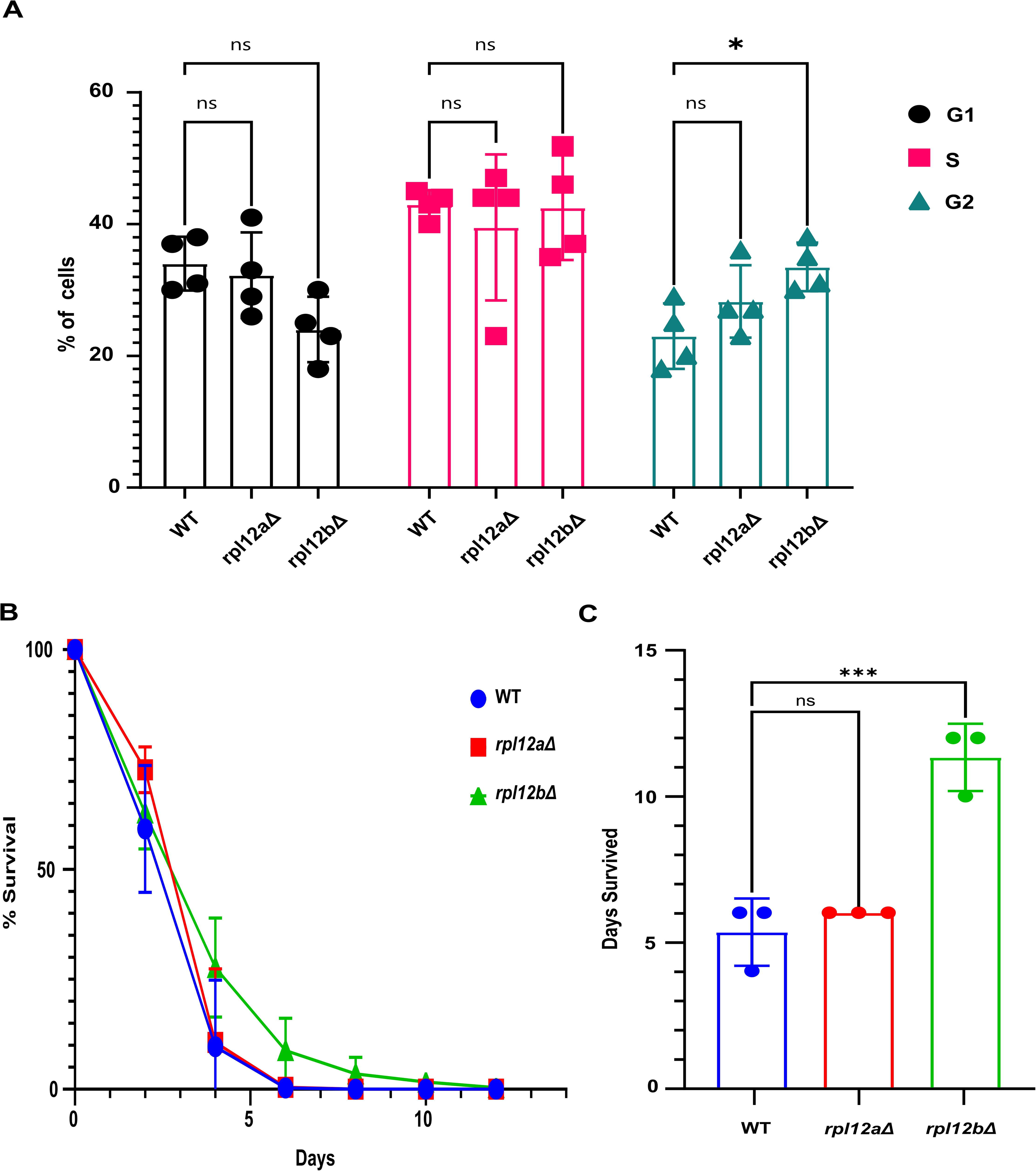
*rpl12aΔ* and *rpl12bΔ* deletion show alters cell cycle progression and differential chronological lifespan. **(A)** Comparative cell cycle analysis showing the percentage of cells in G1, S, and G2 phases for wild-type BY4741, *rpl12aΔ*, and *rpl12bΔ* strains. Cell cycle distribution was analyzed using two-way ANOVA, revealing a significant decrease in G1 population for *rpl12bΔ* compared to wild type (p < 0.05). **(B)** Chronological lifespan analysis showing survival curves of the three strains over time. **(C)** Quantification of lifespan as mean days survived for each strain analysed using ordinary one-way ANOVA with indicated statistical significance according to adjusted P values. Error bars represent SEM.

Chronological lifespan (CLS) analysis further revealed a significant extension of viability in the *rpl12bΔ* strain compared to the WT and the *rpl12aΔ* mutant. *rpl12bΔ* cells remains viable for approximately 10–12 days, whereas WT and *rpl12aΔ* exhibited a shorter lifespan of 5–7 days (**Fig.2 B)**. The statistical analysis confirmed that the extended survival of *rpl12bΔ* is highly significant (p < 0.001), while the *rpl12aΔ* mutant showed no statistically significant difference relative to wild type (**Fig.2 C)**.

Together, these findings point to a paralog-specific role of Rpl12b in regulating cellular lifespan, the G2/M delay observed in *rpl12bΔ* may reduce proliferative activity and promote survival during stationary phase, a phenotype commonly associated with reduced TOR signaling. This suggests the loss of Rpl12b impairs TOR dependent growth programs, leading to adaptive translational and metabolic changes that ultimately extend lifespan.

### Enhanced amino acid biosynthesis and metabolic reprogramming in the *rpl12bΔ*

To validate the differential chronological life span, we performed untargeted metabolic profiling through LC-MS in both positive and negative ionization modes to assess the different metabolic regulations through intracellular metabolites across WT, *rpl12aΔ*, and *rpl12bΔ* strains (**Fig.3 A & B)**. The heatmap clustering of metabolite profile shows that *rpl12bΔ* exhibits a distinct metabolic signature compared to WT and *rpl12aΔ*.

**Figure 3.**
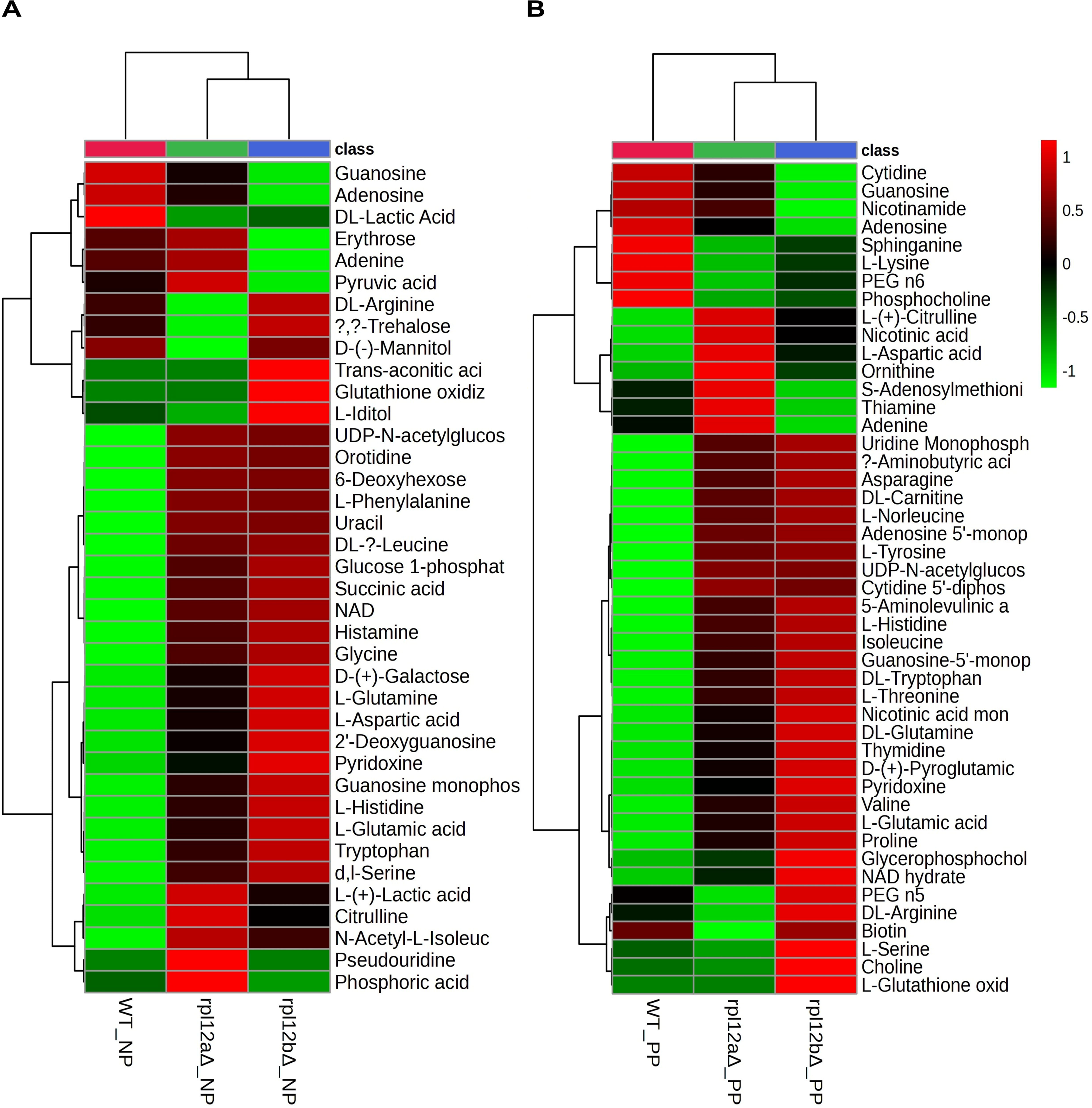
Enhanced amino acid biosynthesis and metabolic reprogramming in the *rpl12bΔ* strains. Heatmaps depict hierarchical clustering of metabolites in WT, *rpl12aΔ*, and *rpl12bΔ* strains under YPD conditions, analyzed in A) negative polarity (NP) and B) positive polarity (PP, right) in YPD conditions. The color scale (−1 to +1) represents normalized metabolite abundance (green = low, red = high), revealing genotype-specific alterations in nucleotide, amino acid, and energy metabolism.

Specifically, the *rpl12bΔ* shows higher levels of several amino acids, glutamate, serine, aspartate, histidine, arginine, tryptophan, leucine, and proline which are generally decreased in the absence of Gcn4-dependent transcriptional programming, suggesting activation of amino acid biosynthetic pathways. In addition, *rpl12bΔ* showed increased abundance of redox-associated metabolites such as NAD and glutathione, consistent with enhanced redox homeostasis.

Metabolites linked to central carbon metabolism were also enriched in *rpl12bΔ*, including glycolytic intermediates (pyruvate, lactate) and TCA cycle-related metabolites, indicating coordinated metabolic remodeling rather than isolated pathway changes. Moreover, the accumulation of stress-related carbohydrates like trehalose and mannitol indicates the presence of stress adaptation responses that are usually linked to longevity. In contrast, *rpl12aΔ* showed relatively mild metabolic changes compared to WT, which is in line with its mild phenotype for lifespan and underscores the metabolic role of Rpl12b.

Together, these findings suggest a model in which loss of Rpl12b drives widespread metabolic reprogramming, including increased amino acid biosynthesis, improved redox status, and activated stress response metabolic pathways. This profile of metabolic reprogramming is consistent with decreased TOR signaling and increased Gcn4 activity, which links ribosomal paralog composition to translational control of metabolic state. Critically, our findings are consistent with previous studies suggesting that decreased levels of certain 60S ribosomal proteins can increase lifespan by modulating translation and promoting metabolic responses associated with amino acid availability and stress response. (Steffen et al. 2008b; Wu et al. 2013; Gulias et al. 2023b).

The findings suggest that loss of Rpl12b specifically induces translational reprogramming and metabolic adaptation through Gcn4 activation, increased amino acid synthesis, and mitochondrial activity to support stress resistance and extended lifespan, whereas loss of *RPL12a* does not induce such adaptive responses.

### Gcn4-dependent translational and metabolic reprogramming underlies longevity in *rpl12bΔ*

To determine whether the observed metabolic changes in *rpl12bΔ* correlated with translational changes, we examined the overlap between differentially translated genes and previously reported Gcn4 target genes (Mittal et al. 2017a). Using published PUNCH-P translational profiling data (Singh et al. 2024), we analyzed the top 50 Gcn4-mediated genes across *rpl12aΔ* and *rpl12bΔ* mutants.

The heatmap (**Fig. 4A**) reveals clear paralog-specific translational regulation, with *rpl12bΔ* showing stronger differential expression of Gcn4-associated metabolic and biosynthetic genes compared with *rpl12aΔ*. Notably, several genes involved in amino acid biosynthesis and metabolic regulation including Lys1, Pck1, Agr7, Arg4, Arg1, Arg6, Rsb1, Rpb4, and Asn1 are known Gcn4 targets strongly upregulated, especially in *rpl12bΔ* contribute to amino acid synthesis. In addition, the genes involved in nitrogen metabolism like Gdh2, Gpd2 and mitochondrial function like Mae1 also elevated in *rpl12bΔ.* However, genes related to protein synthesis and mitochondrial function, such as Rpl28, Sup6, Rps8b, Ssb2, and Grs1, appear more strongly downregulated in *rpl12bΔ*, which depicts severe suppression of cellular and translation-related processes consistent with a shift from growth-associated translation toward stress-adaptive translational programs.

**Figure 4.**
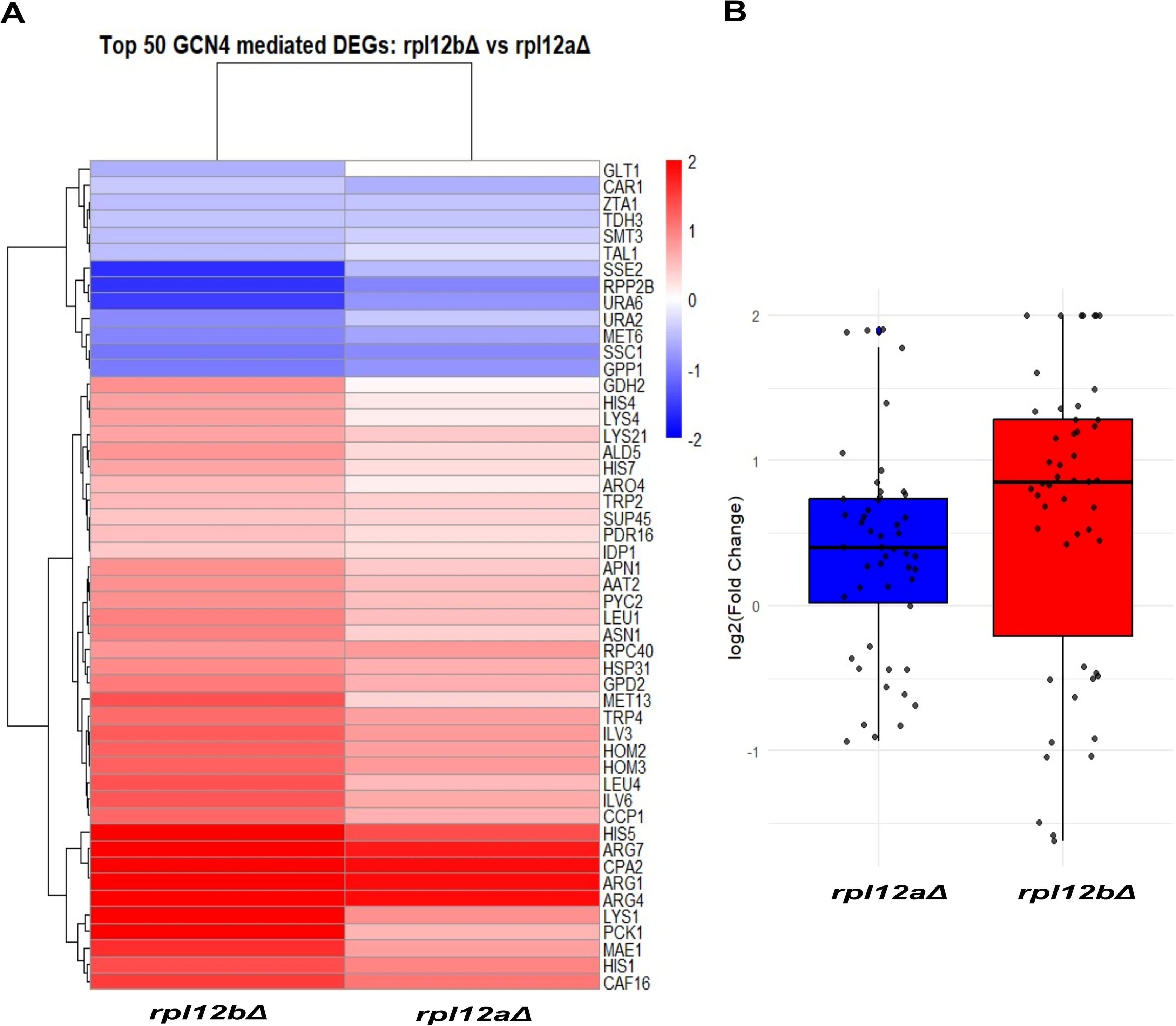
Gcn4-dependent translational and metabolic reprogramming underlies longevity in *rpl12bΔ*. **(A)** Heatmap showing the top 50 Gcn4-mediated differentially expressed genes (DEGs) comparing *rpl12bΔ* and *rpl12aΔ* strains. Gene expression values are represented as log₂ fold change and displayed using a blue, white–red colour scale, where blue indicates downregulation and red indicates upregulation. **(B)** Box-and-whisker plot summarizing the distribution of log₂ fold change values for the selected GCN4 target genes in *rpl12aΔ* and *rpl12bΔ* strains. Individual data points represent gene-level expression changes. Error bars represent SEM.

The box plot (**Fig. 4B**) shows comparative log₂ fold changes between *rpl12aΔ* and *rpl12bΔ* demonstrates a broader distribution and greater variability of gene expression changes in *rpl12bΔ*, supporting the presence of more extensive translational reprogramming, although overall differences are not uniformly significant. Together, these findings suggest that *rpl12bΔ* induces a Gcn4-associated translational program that enhances amino acid biosynthesis while reducing translation-related processes, linking metabolic remodeling with lifespan extension under reduced TOR signaling.(Steffen et al. 2008b; Mittal et al. 2017a).

### Rpl12b maintains TOR-dependent Rps6 phosphorylation without inducing CDS length-dependent translational bias

To investigate whether Rpl12 paralogs differentially influence TOR signaling, we examined phosphorylation of ribosomal protein S6 under nutrient-rich and TOR-inhibited conditions. At the molecular level, immunoblot analysis demonstrated that phosphorylation of ribosomal protein S6 at serine residues 232 and 233 was clearly reduced or absent in *rpl12bΔ* (p < 0.001) under nutrient-rich conditions, while maintained in *rpl12aΔ* (p <0.05) as compared to wild type (**Fig.5 A, B).** Total Rps6 levels were significantly decreased in both *rpl12aΔ* and *rpl12bΔ* mutants (p <0.05), but loading controls remained comparable. Under rapamycin treatment, Rps6 phosphorylation abolished across all strains, confirming TOR inactivation, although a reduced level of total Rps6 protein was observed across all samples. The absence of Rps6 phosphorylation specifically in the *rpl12bΔ* mutant under nutrient-rich conditions suggests a potential Rpl12b-dependent regulation of Rps6 phosphorylation contributes to maintaining TORC1-dependent signaling at the ribosome. Rapamycin treatment for 60 minutes likely results in TOR inactivation, leading to dephosphorylation at conserved serine residues. Additionally, the cellular stress induced by rapamycin may activate ribophagy, contributing to the overall reduction in Rps6 protein levels.

**Figure 5.**
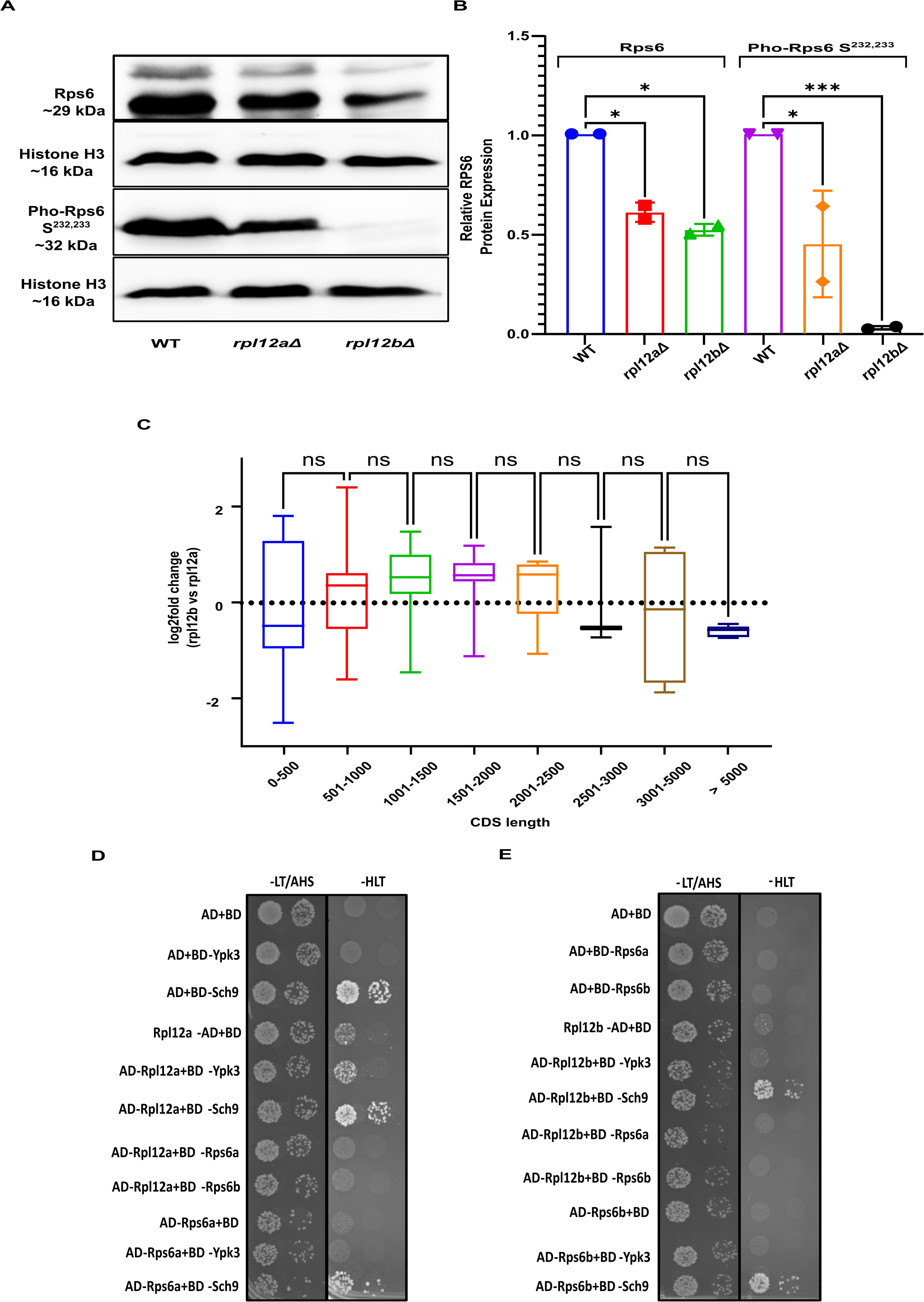
Rpl12b maintains TOR-dependent Rps6 phosphorylation without inducing CDS length-dependent translational bias. **(A)** Western blot analysis of total Rps6 and phospho-Rps6 levels in *rpl12aΔ* and *rpl12bΔ* strains relative to wild type (WT). Histone H3 was used as a loading control. **(B**) Quantification of total Rps6 and phospho-Rps6 protein levels normalized to WT. **(C)** Phospho-Rps6 occupancy analysis comparing *rpl12bΔ* and *rpl12aΔ* across transcripts grouped by coding sequence (CDS) length bins. **(D, E)** Yeast two-hybrid (Y2H) assays were performed to assess potential direct interactions between TOR regulators and Rpl12 paralogs. Error bars represent SEM.

Because Rps6 phosphorylation has been associated with enhanced translation of mRNAs with short coding sequences (CDSs), with progressively reduced phosphorylation observed on ribosomes translating longer CDSs suggests the increase in translational efficiency for short CDS mRNAs and establishes a length-dependent regulatory mechanism mediated by Rps6 phosphorylation (Bohlen et al. 2021). To investigate whether a similar CDS length dependent translation bias exists between the yeast ribosomal protein paralogs *RPL12*a and *RPL12*b we analyzed previously available Punch-P data (Singh et al. 2024), using log₂ (fold change) in gene expression between *rpl12aΔ* and *rpl12bΔ* strains (*rpl12aΔ* vs *rpl12bΔ*) across CDS length in YPD medium. No statistically significant differences (**Fig. 5C**) were observed across CDS length categories (0–500 nt to >5000 nt), as indicated by the absence of CDS-dependent bias in the box plot analysis. Unlike Rps6 phosphorylation, *RPL12* paralog-specific translation does not exhibit CDS length-dependent bias, suggesting that their functional specialization may operate through mechanisms unrelated to elongation dependent modification. Notably, western blot analysis confirmed the absence of detectable Rps6 phosphorylation in the *rpl12bΔ* strain under YPD conditions, implying that Rpl12b is required for maintaining Rps6 phosphorylation, at least in rich medium. However, the *rpl12bΔ* mutant, which lacks CDS-lengthdependent translation changes, indicates that its deletion does not replicate the translational phenotype observed in Rps6 phospho-deficient cells. This suggests potential mechanistic variation in yeast, possibly involving compensatory pathways or alternative modes of ribosomal specialization distinct from the regulatory mechanisms seen in mammalian systems.

To determine whether this phenotypic regulation might arise from direct physical interactions, we performed Yeast2hybrid to check protein–protein interaction between Rpl12a/b and TORC1 effectors including Rps6 paralogs, Ypk3, and Sch9. No direct associations were detected (**Fig. 5D, E**), indicating that *Rpl12* paralogs likely modulate TOR signaling indirectly, through altered ribosomal stalk composition and downstream regulatory pathways rather than physical association.

### Paralog dependent expression of Ribosome preservation factor Stm1

To determine whether ribosome preservation factors contribute to the phenotypes observed in *RPL12* paralog mutants, we examined Stm1 at both transcript and protein levels to see if it contributes to the differences between the paralog mutants. qPCR analysis revealed no significant differences in *STM1* mRNA across wild type, *rpl12aΔ*, and *rpl12bΔ* strains (**Fig. 6A**), indicating *STM1* is not transcriptionally regulated by RPL12 paralogs.

**Figure 6.**
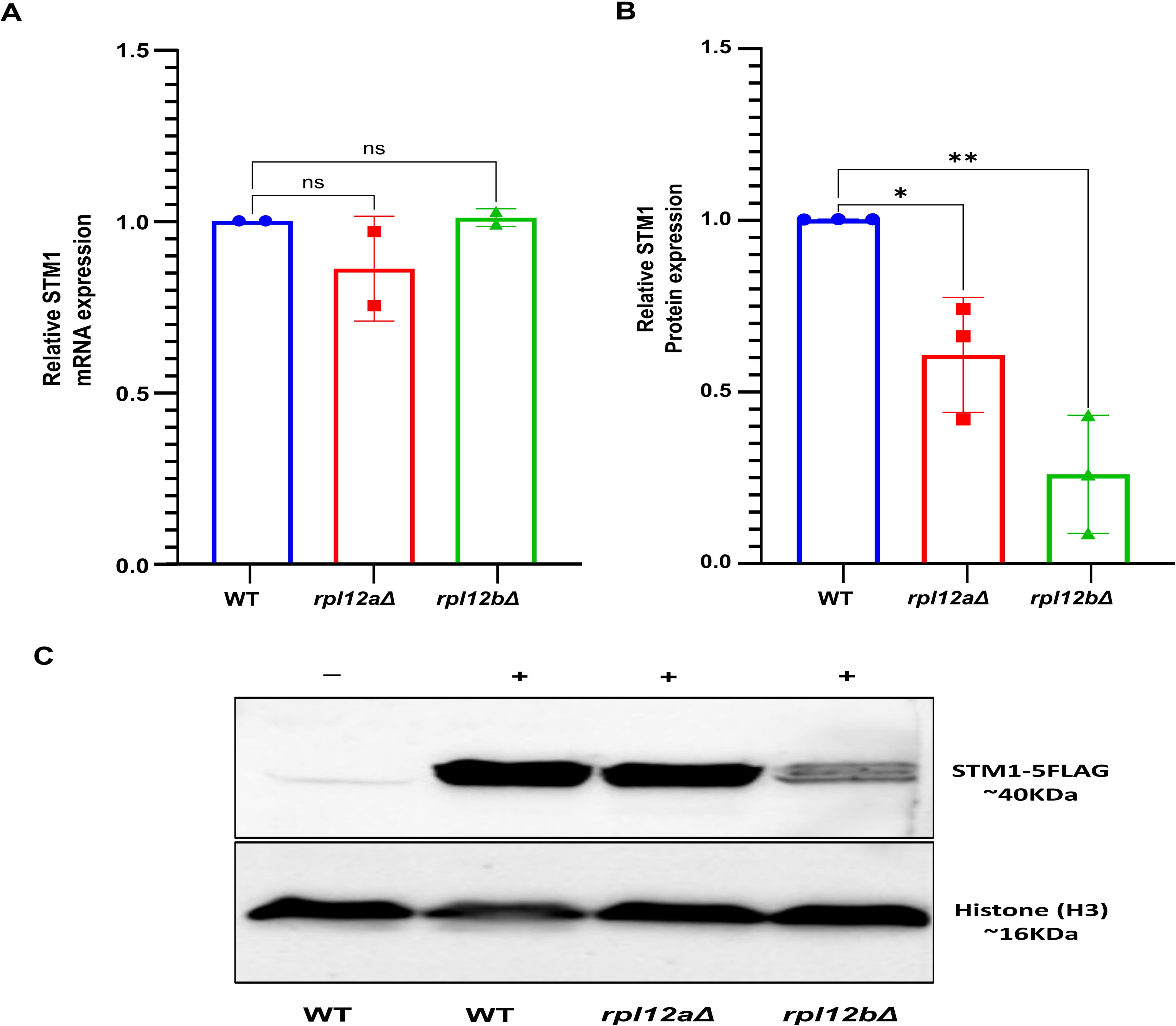
Paralog dependent expression of Ribosome preservation factor Stm1. **(A)** Relative *STM1* mRNA levels in *rpl12aΔ* and *rpl12bΔ* strains compared to wild type (WT), quantified by RT–qPCR in YPD medium and normalized to *HHF1* mRNA. **(B)** Quantification of Stm1 protein levels from total cell lysates expressing C-terminally Flag-tagged Stm1 in rpl12aΔ and rpl12bΔ strains relative to wild type (WT). **(C)** Representative Western blot showing Flag-tagged Stm1, with Histone H3 serving as the loading control. Error bars represent SEM.

To assess protein-level regulation, Stm1 was endogenously FLAG-tagged at the C-terminus in wild type and paralog deletion strains: *rpl12aΔ*, and *rpl12bΔ* strains. Immunoblot analysis revealed a modest reduction of Stm1 protein in *rpl12aΔ* (p <0.05) and a pronounced decrease in *rpl12bΔ* (p < 0.01) under nutrient-rich conditions (**Fig. 6B,C**). These findings indicate that loss of *RPL12b* affects Stm1 primarily at the post-transcriptional level. Because Stm1 functions as a ribosome preservation factor that stabilizes inactive 80S ribosomes during stress, its diminished protein levels suggest impaired ribosome preservation capacity in the absence of *RPL12*b.

Together, these results indicate that Rpl12b influences Stm1 protein abundance and thereby contributes to ribosome preservation. This supports a model in which ribosomal stalk composition regulates ribosome stability through Stm1, linking paralog-specific ribosome architecture to translational control and cellular stress adaptation.

## DISCUSSION

Ribosomal paralogs are increasingly recognized as functional regulators rather than redundant structural components. In this study, we demonstrate that the ribosomal stalk paralogs *RPL12a* and *RPL12b* in *S. cerevisiae* perform specialized, non-redundant functions that become particularly evident under TOR pathway perturbation. While both *rpl12aΔ* and *rpl12bΔ* strains showed only modest growth defects in rich medium, rapamycin treatment unmasked striking paralog-specific phenotypes. Deletion of *RPL12a* enhanced growth, whereas deletion of *RPL12b* severely impaired proliferation. The paralog specific effect under rapamycin conditions, despite successful rescue in nutrient-rich medium, underscores that *RPL12a* and *RPL12b* confer specialized roles in translational regulation rather than acting as interchangeable structural components that extend beyond simple redundancy (Malik Ghulam et al. 2022; Komili et al. 2007).

Expression analyses indicate hierarchical regulation between the paralogs, we found that *RPL12b* significantly expressed at higher level than *RPL12a* in wildtype, under both normal and TOR inhibition, indicating a clear transcriptional bias. This trend is also reflected at the protein level, where Rpl12b shows slightly higher abundance, suggesting that the increased expression is maintained from mRNA to protein. This suggests that *RPL12*b functions as the dominant paralog in wild type cells, maintaining ribosomal stoichiometry.

At the physiological level, *rpl12bΔ* displayed altered cell-cycle distribution with increased G2/M accumulation together with a pronounced extension of chronological lifespan. This phenotype is consistent with reduced growth signaling and resembles longevity observed under nutrient limitation or reduced ribosome function, where cells shift toward stress resistance and maintenance rather than proliferation(Steffen et al. 2008b).

Multi-omics analyses further demonstrated that Rpl12 paralog coordinates a multilayered adaptive program. Loss of *RPL12b* selectively activated Gcn4-dependent translation even in nutrient-rich conditions, mimicking nutrient limitation. This translational reprogramming was accompanied by disrupted CDS length–dependent translation patterns, consistent with altered ribosome mRNA interactions. Metabolomic profiling reinforced this connection, revealing changes in amino acid pools, TCA cycle intermediates, and redox-associated metabolites that align with Gcn4-driven metabolic programs. Together, these data suggest that *RPL12b* deletion induces a Gcn4-centered translational and metabolic response, mirroring reduced TOR activity, and promoting stress tolerance over rapid growth (Steffen et al. 2008b; Mittal et al. 2017a).

Mechanistically, Rps6 phosphorylation analysis provide a direct link of *RPL12*b to TOR signaling. Loss of *RPL12b* reduced phosphorylation of Rps6 at serine 232/233, a canonical TOR readout, indicating Rpl12b as a critical mediator of TOR dependent translation (Yerlikaya et al. 2016). While rapamycin treatment eliminated Rps6 phosphorylation across all strains, the differential expression patterns observed suggest that Rpl12b contributes to maintaining ribosomal integrity under TOR inhibition, potentially through interactions with TOR kinase complexes or downstream effectors. Though Yeast2hybrid, the absence of direct interaction of *RPL12* paralogs with TOR effectors suggests that this regulation occurs indirectly, likely through changes in ribosomal stalk composition that influence downstream signaling.

In addition to Rps6, our data show that Stm1 protein levels are reduced in *rpl12bΔ* despite unchanged mRNA. Because Stm1 is a TORC1-regulated ribosome preservation factor, this reduction further supports the idea that *RPL12b* deletion might destabilizes ribosomes and weakens TORC1-mediated stress protection. While Stm1 is not the primary driver of the phenotypes we observed, its diminished abundance suggests ribosomal stalk protein paralog (Rpl12a or Rpl12b) might indirectly influence ribosome stability and can determine how cells grow, adapt, and live longer.

Together, these findings establish Rpl12b as a novel interface between ribosomal heterogeneity and TOR signaling. By coupling ribosomal stalk composition to nutrient-responsive pathways, Rpl12b deletion reprograms translation and metabolism to favor survival over proliferation. This work underscores the broader principle that ribosomal paralog specialization provides an additional layer of TOR pathway regulation, highlighting ribosome composition as a determinant of cellular adaptation and longevity. Future studies should map the precise structural changes in ribosomes caused by Rpl12b loss, test conservation across conditions and species, and dissect how altered ribosome architecture feeds into specific signaling and translational outcomes.

In summary, *RPL12a* and *RPL12b* are functionally distinct paralogs: Rpl12a primarily supports ribosome homeostasis, whereas Rpl12b links ribosome composition to nutrient signaling. Deletion of *RPL12b* reduces TORC1-dependent Rps6 phosphorylation, causes G2/M cell-cycle perturbation, and extends chronological lifespan, accompanied by a Gcn4-associated translational program and metabolic remodeling (increased amino-acid biosynthesis, altered redox and TCA intermediates). Together, these data support a model in which loss of *RPL12b* shifts cells from growth toward a stress-adaptive metabolic state that favors survival over proliferation, underscoring ribosomal paralog specialization as an additional layer of TOR regulation.

## METHODS

### Yeast Strains Media

All *S. cerevisiae* strains used in this study were derived from the BY4741 (MATa his3Δ1 leu2Δ0 met15Δ0 ura3Δ0); gene deletions and epitope tagging were performed by homologous recombination using PCR-based methods. Yeast cultures were grown in YPD (1% Yeast extract, 2% Peptone, 2% Dextrose) medium or synthetic complete medium (SC), with or without selection at 30°C with continuous shaking (180 rpm) unless otherwise specified. For rapamycin sensitivity assays, cultures were grown in YPD supplemented with 0.2 µg/mL rapamycin (prepared in DMSO) or YPD agar plates containing 0.1 µg/mL rapamycin.

### Growth tests on liquid and solid media

For liquid growth assay, yeast strains were pre-cultured overnight in YPD medium at 30 °C with shaking at 180 rpm. The following day, OD₆₀₀ was measured using a spectrophotometer, and cultures were diluted to a starting OD₆₀₀ of 0.01 in fresh medium with and without rapamycin. The 200 µL culture in triplicates were dispensed into 96-well microplate and subjected to microplate reader (Agilent BioTek Cytation 5) for growth kinetics at 30 °C with shaking and OD₆₀₀ was measured at 30 min intervals pooled into one dataset and analyzed via the Growthcurver R package(Sprouffske and Wagner 2016) and GraphPad.

For the drop test, mid log phage cells normalized with 0.5 OD₆₀₀ and serially diluted 10^-1^ to 10^-5^ 5ul cells spotted on YPD and YPD Rapamycin plate and incubated at 30 °C and photo documented after 2-4 days.

### Cross complementation Assay

The *RPL12a* and *RPL12b* genes, including their native promoter and terminator regions, were PCR-amplified from genomic DNA using primers containing ClaI and SacI restriction sites and cloned into the pRS316 vector. Cross complementation was tested by transforming pRS316-*RPL12b* and pRS316-*RPL12a* in *rpl12aΔ* and *rpl12bΔ* respectively by standard PEG-LiAc(Longtine et al. 1998), plated on selective plate and grown for 2-3 days and subsequently subjected to both YPD and rapamycin conditions to assess functional rescue.

### Chronological Lifespan (CLS) and Cell Cycle Assays

Yeast strains were aged in SC medium containing all amino acids for 72 h at 30°C. Viability was assessed by dilutions on SC-All agar and counting colony-forming units (CFUs) every 48 h. In parallel, cell death was quantified by propidium iodide (PI) staining followed by flow cytometry, providing an independent measure of viability.

For cell cycle analysis, mid-log phase cultures (OD₆₀₀ = 0.5–0.6) were harvested and fixed in 70% ethanol overnight at 4 °C. Fixed cells were washed, treated sequentially with RNase A (1 mg/mL, 1 h, 37 °C) and Proteinase K (1 mg/mL, 1 h, 37 °C), and stained with PI (50 µg/mL). DNA content was analyzed using a BD Accuri C6 flow cytometer (FL-2 PE channel, 488 nm excitation), and cell cycle profiles were generated with FlowJo software to determine the distribution of cells across G1, S, and G2/M phases.

### Yeast two-hybrid assay

The yeast two-hybrid experiment was performed as per the manufacturer’s protocol for the Matchmaker GAL4-based two-hybrid system (Clontech). Equal amounts of AD constructs were mixed with the corresponding BD constructs in separate reactions and the mixtures were introduced into the Y2H Gold yeast strain. The transformed yeast cells were selected on Leu−/Trp−/His− and also on Leu−/Trp−/His−/Ade− in order to screen for stronger interactions. Aliquots plated on Leu−/Trp− medium were used as transformation control. Simultaneously, the cells were diluted 1:10 and 1:100 fold using 1X TE and plated on Leu−/Trp− and Leu−/Trp−/His− medium. The plates were incubated for 3–4 d at 30°C and then photographed.

### C-terminus epitope tagging

RPL12a and RPL12b paralogs were FLAG-tagged at C-terminus using 40bp homologous region before and after the stop codon, followed by the epitope sequence with the stop codon and HIS3 gene as a selection marker extracted via Cre-mediated recombination (Haim-Vilmovsky and Gerst 2009). Following recombination procedure tagged colonies were patch on YPD and SC-HIS plate, and those who fails to grow on SC-HIS subjected to check correct tagging by colony PCR (cPCR) using genomic DNA to confirm correct integration of the FLAG tag.

Stm1 were also FLAG-tagged at C-terminus using 40bp homologous region before and after the stop codon, followed by the epitope sequence with the stop codon and hygromycin B as selection marker in wild type, *rpl12aΔ* and *rpl12bΔ,* correct tagging was verified by PCR on genomic DNA.

### RNA extraction, cDNA synthesis and quantitative real time-PCR (qRT-PCR)

Total RNA was isolated from yeast cultures grown to mid-log phase (OD₆₀₀ = 0.5–0.6). Cells from 10 mL cultures were harvested by centrifugation, washed twice with sterile PBS, and lysed using a formamide–EDTA protocol (Shedlovskiy et al. 2017) followed by DNaseI (NEB) treatment for 1 hour. For cDNA synthesis, 1.5 µg of total RNA was reverse-transcribed using the High-Capacity cDNA Reverse Transcription Kit (Applied Biosystems) following the manufacturer’s instructions. Quantitative real time PCR (RT-PCR) was performed using iTaq Universal SYBR Green Supermix (BioRad) using gene-specific primers listed in Table 2. Reactions were performed in triplicate with three biological replicates under the following cycling conditions: initial denaturation at 95 °C for 10 min, followed by 40 cycles of 95 °C for 15 s and 60 °C for 1 min. The specificity of individual real-time PCR products was assessed by melting-curve analysis. Melting curves for individual PCR products displayed a single peak. Expression levels were normalized to HHF1 as the internal reference gene, and relative fold changes were calculated using the ΔΔCt method. Statistical significance was determined using ordinary one-way Anova, with p < 0.05 considered significant.

### Total Protein Extraction

Yeast cells were cultured to mid-log phase (OD_600_ = 0.5–0.6) and treated with 0.2ug/ml rapamycin for 1 hour then harvested for protein extraction. For phosphorylation-sensitive analyses, cells were resuspended in Tris-based lysis buffer [(50 mM Tris-HCl pH 7.5, 150 mM NaCl, 15% glycerol, 0.5% Tween-20) supplemented with a phosphatase inhibitor mixture (10 mM NaF, 10 mM NaN3, 10 mM p-nitrophenylphosphate, 10 mM sodium pyrophosphate, and 10 mM β-glycerophosphate) and an EDTA-free protease inhibitor cocktail (Roche)]. In parallel, cells were also lysed in RIPA buffer [50 mM Tris-HCl pH 7.4, 150 mM NaCl, 1% Triton x-100, 0.5% sodium deoxycholate, 0.1% SDS, and an EDTA-free protease inhibitor cocktail (Roche)] for total protein extraction. Cells were lysed by bead beating (0.5 mm beads, 30 Hz, three 15-minute cycles) and lysate were harvested by centrifugation at 13,000 g for 10 min at 4°C to remove cell debris. Protein concentrations were determined using Bradford assay.

### Western Blotting

Total cell lysates and ribosome fractions were separated by SDS-PAGE and the proteins were transferred on to nitrocellulose membrane (0.45 μm pore size; BioRad). Membranes were blocked with 5% BSA for 1 hour at room temperature incubated overnight at 4 °C with primary antibodies (1:2000 dilution) against Rpl12 (Invitrogen), phospho-Rps6 (Ser235/236), Rps6, and Histone H3 (Cell Signaling). After washing, membranes were incubated with HRP-conjugated anti-rabbit or anti-mouse IgG secondary antibodies (1:5000 dilution; Invitrogen) for 1 h at room temperature. Signals were detected using Clarity Western ECL substrate (Bio-Rad) and imaged with a Chemidoc system (Bio-Rad). Finally, the band intensity was measured using ImageLab or ImageJ software, and statistical plots were generated in GraphPad Prism.

### Metabolite Extraction from Yeast Cells

Metabolite extraction method followed by previously described in Mohammad et. al. 2021 (Mohammad et al. 2021) with minor modifications. Yeast cultures were grown for 24 h in triplicate, and 10 ml aliquots (∼5.0 × 10⁸ cells) 50-ml Falcon tubes. The total volume was adjusted to 40 ml with prechilled (−20°C) quenching solution (60% methanol in 155 mM ABC buffer, pH 8.0). Samples were centrifuged (3 min, 11,325 × g, −5°C; MPW-150R Refrigerated Centrifuge), and the supernatant was removed carefully without disturbing the pellet. The pellet was resuspended in 10 ml of ice-cold ABC buffer, transferred to 15-ml centrifuge tubes, and pelleted again (3 min, 3,000 × g, 0°C). The quenched pellets were kept ice and processed simultaneously or at −80°C until metabolite extraction.

For extraction, each sample was supplemented with 2 ml of chloroform and 1 ml of methanol (both prechilled to −20°C), 1 ml of ice-cold nanopure water, and 200 μl of acid-washed glass beads (425–600μm). Tubes mouths were sealed with aluminum foil and vortexed for 30 min at 4°C to facilitate extraction. Samples were then incubated on ice for 30 min to allow protein precipitation and phase separation, followed by centrifugation (10 min, 3,000 × g, 4°C). The ∼400 μl upper aqueous phase (containing water-soluble metabolites) was transferred to a 1.5-ml Eppendorf tube containing 800 μl of acetonitrile prechilled to −20°C. After centrifugation (10 min, 13,400 × g, 4°C), ∼800 μl of the clarified supernatant was transferred to LC–MS vials and stored at 0°C prior to LC–MS/MS analysis.

The extracted water-soluble metabolites from yeast samples and analyzed by Liquid Chromatography run was performed using UHPLC (Thermo Scientific Vanquish, MA, USA) coupled to an Orbitrap (Exploris 240 HRMS, Thermo Fisher Scientific, MA, USA) equipped with a heated electrospray ion source (H-ESI). The separation and detection of metabolites achieved using a syncronis HILIC column (Thermo Scientific; 150 mm x 2.1 mm, 5 μm particle size)

Data processing was performed using Compound Discoverer v3.3 software (Thermo Fisher Scientific) with an m/z range of 70-1000. Metabolite identification was based on MS² peak analysis, while quantification was carried out using MS¹ peak intensities within the software workflow. Processed data were subsequently visualized and plotted using the freely available tool MetaboAnalyst 4.0 (Pang et al. 2024).

### Material availability

Yeast strains used in this study (Supplementary Table X)

## Acknowledgements

The author thanks Dr. Jeffrey E. Gerst (Weizmann Institute of Science, Israel) for BY4741, rpl12aΔ, and rpl12bΔ. We thank Ms. Meghavi Raval for the STM1-5X Flag, Dr. Krishna Swamy (Ahmedabad University) for the *tor1Δ* strain, pUG27 loxP-pAgTEF1-Sphis5-tAgTEF1-loxP, and pSH47 plasmids, and Dr. Vivek Verma (Gujarat Biotechnology University) for the Y2HGold strain, pGADT7, and pGBKT7 plasmids, Dr. Gaurav Jerarth (Gujarat Biotechnology University) for bioinformatics help. The authors would like to thank Gujarat Biotechnology University, Gandhinagar, Gujarat, India, for providing a space to work and instrumentation facilities to complete this work. This work was supported by GBU core grant and DBT-RLS grant.

---

Yeast Strains

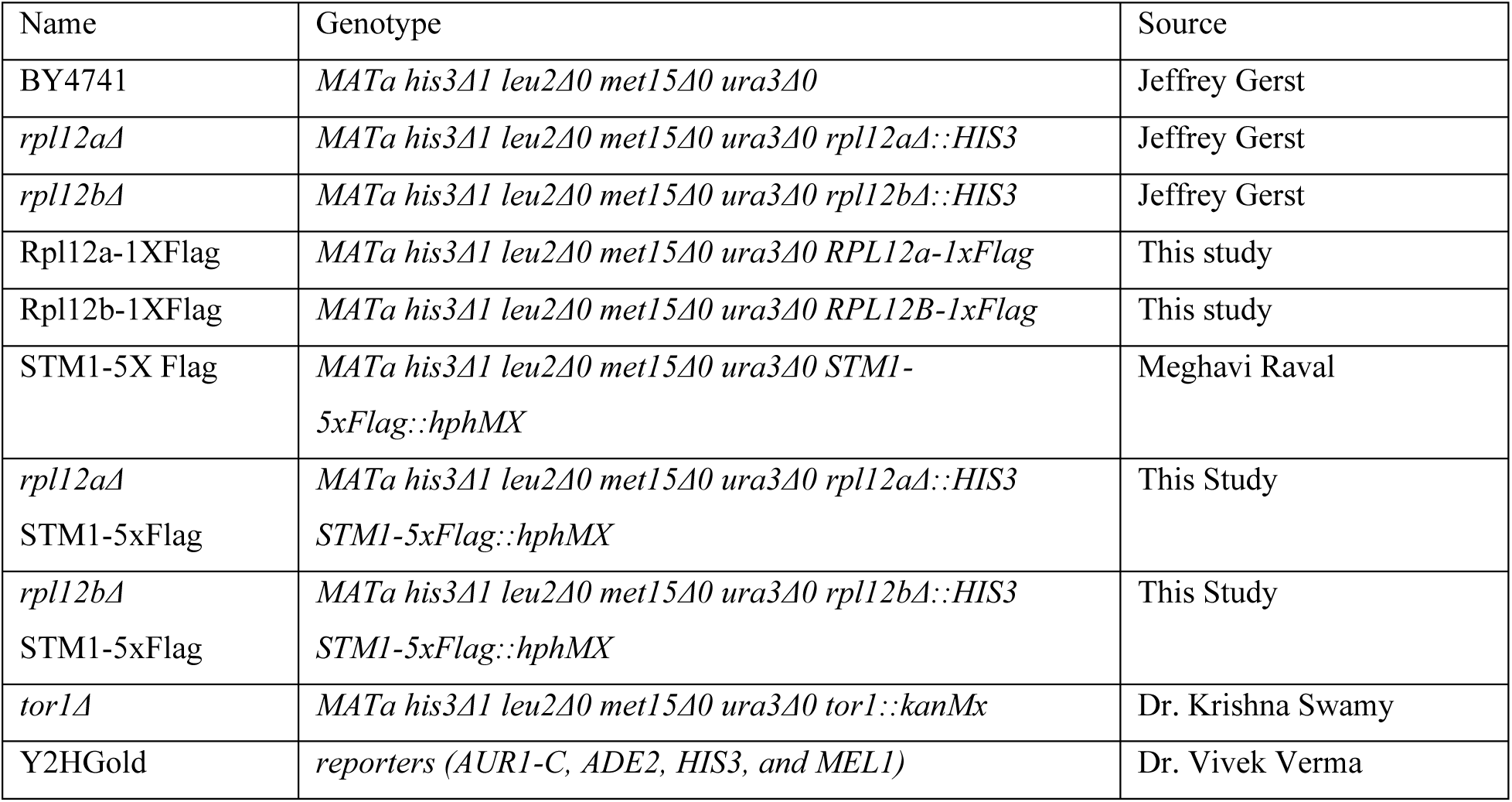

Plasmids

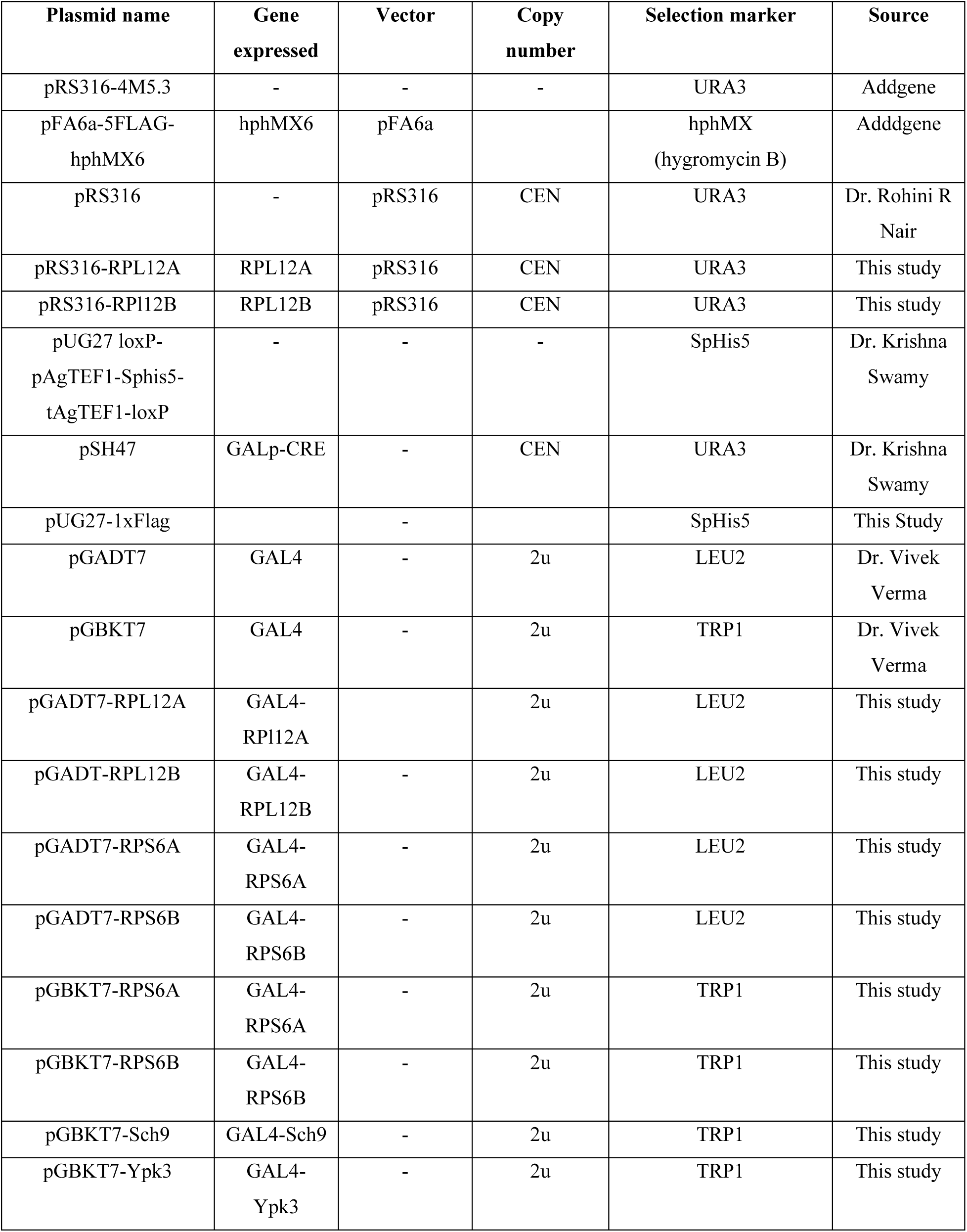

## REFERENCES

Aspden, Julie, William James Faller, Maria Barna, and Anders Lund. 2025. “Ribosome Heterogeneity and Specialization.” Philosophical Transactions of the Royal Society of London. Series B, Biological Sciences 380 (1921): 20230375. 10.1098/rstb.2023.0375.

Bohlen, Jonathan, Mykola Roiuk, and Aurelio A. Teleman. 2021. “Phosphorylation of Ribosomal Protein S6 Differentially Affects mRNA Translation Based on ORF Length.” Nucleic Acids Research 49 (22): 13062–74. 10.1093/nar/gkab1157.

Briones, Elisa, Carlos Briones, Miguel Remacha, and Juan P. G. Ballesta. 1998. “The GTPase Center Protein L12 Is Required for Correct Ribosomal Stalk Assembly but Not for Saccharomyces cerevisiaeViability *.” Journal of Biological Chemistry 273 (48): 31956–61. 10.1074/jbc.273.48.31956.

Chen, Yuting, Jiaxin Hu, Pengwei Zhao, et al. 2025. “Rpl12 Is a Conserved Ribophagy Receptor.” Nature Cell Biology 27 (3): 477–92. 10.1038/s41556-024-01598-2.

Crick, F. H. 1958. “On Protein Synthesis.” Symposia of the Society for Experimental Biology 12: 138–63.

Deprez, Marie-Anne, Elja Eskes, Joris Winderickx, and Tobias Wilms. 2018. “The TORC1-Sch9 Pathway as a Crucial Mediator of Chronological Lifespan in the Yeast Saccharomyces Cerevisiae.” FEMS Yeast Research 18 (5). 10.1093/femsyr/foy048.

García-Marcos, Alberto, Antonio Morreale, Esther Guarinos, et al. 2007. “In Vivo Assembling of Bacterial Ribosomal Protein L11 into Yeast Ribosomes Makes the Particles Sensitive to the Prokaryotic Specific Antibiotic Thiostrepton.” Nucleic Acids Research 35 (21): 7109–17. 10.1093/nar/gkm773.

Genuth, Naomi R., and Maria Barna. 2018. “The Discovery of Ribosome Heterogeneity and Its Implications for Gene Regulation and Organismal Life.” Molecular Cell 71 (3): 364–74. 10.1016/j.molcel.2018.07.018.

González, Asier, and Michael N. Hall. 2017. “Nutrient Sensing and TOR Signaling in Yeast and Mammals.” The EMBO Journal 36 (4): 397–408. 10.15252/embj.201696010.

Gulias, Juan Facundo, Florencia Niesi, Martín Arán, Susana Correa-García, and Mariana Bermúdez-Moretti. 2023a. “Gcn4 Impacts Metabolic Fluxes to Promote Yeast Chronological Lifespan.” PloS One 18 (10): e0292949. 10.1371/journal.pone.0292949.

Haim-Vilmovsky, Liora, and Jeffrey E. Gerst. 2009. “M-TAG: A PCR-Based Genomic Integration Method to Visualize the Localization of Specific Endogenous mRNAs in Vivo in Yeast.” Nature Protocols 4 (9): 1274–84. 10.1038/nprot.2009.115.

Ikai, Nobuyasu, Norihiko Nakazawa, Takeshi Hayashi, and Mitsuhiro Yanagida. 2011. “The Reverse, but Coordinated, Roles of Tor2 (TORC1) and Tor1 (TORC2) Kinases for Growth, Cell Cycle and Separase-Mediated Mitosis in Schizosaccharomyces Pombe.” Open Biology 1 (3): 110007. 10.1098/rsob.110007.

Imami, Koshi, Miha Milek, Boris Bogdanow, et al. 2018. “Phosphorylation of the Ribosomal Protein RPL12/uL11 Affects Translation during Mitosis.” Molecular Cell 72 (1): 84–98.e9. 10.1016/j.molcel.2018.08.019.

Jacinto, Estela, and Michael N. Hall. 2003. “Tor Signalling in Bugs, Brain and Brawn.” Nature Reviews. Molecular Cell Biology 4 (2): 117–26. 10.1038/nrm1018.

Komili, Suzanne, Natalie G. Farny, Frederick P. Roth, and Pamela A. Silver. 2007. “Functional Specificity among Ribosomal Proteins Regulates Gene Expression.” Cell 131 (3): 557–71. 10.1016/j.cell.2007.08.037.

Li, Dan, and Jianlong Wang. 2020. “Ribosome Heterogeneity in Stem Cells and Development.” Journal of Cell Biology 219 (6): e202001108. 10.1083/jcb.202001108.

Loewith, Robbie, Estela Jacinto, Stephan Wullschleger, et al. 2002. “Two TOR Complexes, Only One of Which Is Rapamycin Sensitive, Have Distinct Roles in Cell Growth Control.” Molecular Cell 10 (3): 457–68. 10.1016/S1097-2765(02)00636-6.

Longtine, M. S., A. McKenzie, D. J. Demarini, et al. 1998. “Additional Modules for Versatile and Economical PCR-Based Gene Deletion and Modification in Saccharomyces Cerevisiae.” *Yeast (Chichester*, England) 14 (10): 953–61. 10.1002/(SICI)1097-0061(199807)14:10%253C953::AID-YEA293%253E3.0.CO;2-U.

Mager, W. H., R. J. Planta, J. P. G. Ballesta, et al. 1997. “A New Nomenclature for the Cytoplasmic Ribosomal Proteins of Saccharomyces Cerevisiae.” Nucleic Acids Research 25 (24): 4872–75. 10.1093/nar/25.24.4872.

Malik Ghulam, Mustafa, Mathieu Catala, Gaspard Reulet, Michelle S. Scott, and Sherif Abou Elela. 2022. “Duplicated Ribosomal Protein Paralogs Promote Alternative Translation and Drug Resistance.” Nature Communications 13 (1): 4938. 10.1038/s41467-022-32717-y.

McIntosh, Kerri B., and Jonathan R. Warner. 2007. “Yeast Ribosomes: Variety Is the Spice of Life.” Cell 131 (3): 450–51. 10.1016/j.cell.2007.10.028.

Melnikov, Sergey, Adam Ben-Shem, Nicolas Garreau de Loubresse, Lasse Jenner, Gulnara Yusupova, and Marat Yusupov. 2012. “One Core, Two Shells: Bacterial and Eukaryotic Ribosomes.” Nature Structural & Molecular Biology 19 (6): 560–67. 10.1038/nsmb.2313.

Mittal, Nitish, Joao C. Guimaraes, Thomas Gross, et al. 2017a. “The Gcn4 Transcription Factor Reduces Protein Synthesis Capacity and Extends Yeast Lifespan.” Nature Communications 8 (1): 457. 10.1038/s41467-017-00539-y.

Mohammad, Karamat, Heng Jiang, Vladimir I. Titorenko, and Karamat Mohammad. 2021. “Quantitative Metabolomics of Saccharomyces Cerevisiae Using Liquid Chromatography Coupled with Tandem Mass Spectrometry.” Journal of Visualized Experiments (JoVE), no. 167 (January): e62061. 10.3791/62061.

Palade, G. E. 1955. “A Small Particulate Component of the Cytoplasm.” The Journal of Biophysical and Biochemical Cytology 1 (1): 59–68. 10.1083/jcb.1.1.59.

Pang, Zhiqiang, Lei Xu, Charles Viau, et al. 2024. “MetaboAnalystR 4.0: A Unified LC-MS Workflow for Global Metabolomics.” Nature Communications 15 (1): 3675. 10.1038/s41467-024-48009-6.

Powers, R. Wilson, Matt Kaeberlein, Seth D. Caldwell, Brian K. Kennedy, and Stanley Fields. 2006. “Extension of Chronological Life Span in Yeast by Decreased TOR Pathway Signaling.” Genes & Development 20 (2): 174–84. 10.1101/gad.1381406.

Powers, Ted, and Peter Walter. 1999. “Regulation of Ribosome Biogenesis by the Rapamycin-Sensitive TOR-Signaling Pathway in Saccharomyces Cerevisiae.” Molecular Biology of the Cell 10 (4): 987–1000. 10.1091/mbc.10.4.987.

Rohde, John R., Robert Bastidas, Rekha Puria, and Maria E. Cardenas. 2008. “Nutritional Control via Tor Signaling in Saccharomyces Cerevisiae.” Current Opinion in Microbiology 11 (2): 153–60. 10.1016/j.mib.2008.02.013.

Sauert, Martina, Hannes Temmel, and Isabella Moll. 2015. “Heterogeneity of the Translational Machinery: Variations on a Common Theme.” *Biochimie*, Quality Control in Protein Synthesis, vol. 114 (July): 39–47. 10.1016/j.biochi.2014.12.011.

Shedlovskiy, Daniel, Natalia Shcherbik, and Dimitri G. Pestov. 2017. “One-Step Hot Formamide Extraction of RNA from Saccharomyces Cerevisiae.” RNA Biology 14 (12): 1722–26. 10.1080/15476286.2017.1345417.

Shetty, Sunil, Jon Hofstetter, Stefania Battaglioni, Danilo Ritz, and Michael N. Hall. 2023. “TORC1 Phosphorylates and Inhibits the Ribosome Preservation Factor Stm1 to Activate Dormant Ribosomes.” The EMBO Journal 42 (5): EMBJ2022112344. 10.15252/embj.2022112344.

Singh, Raman K., Robert A. Crawford, Dheerendra P. Mall, Graham D. Pavitt, and Jeffrey E. Gerst. 2024. “The 3’-Untranslated Regions of Yeast Ribosomal Protein mRNAs Determine Paralog Incorporation into Ribosomes and Recruit Factors Necessary for Specialized Functions.” Preprint, bioRxiv, March 19. 10.1101/2024.03.18.585503.

Sprouffske, Kathleen, and Andreas Wagner. 2016. “Growthcurver: An R Package for Obtaining Interpretable Metrics from Microbial Growth Curves.” BMC Bioinformatics 17 (1): 172. 10.1186/s12859-016-1016-7.

Stark, Holger, Marina V. Rodnina, Jutta Rinke-Appel, Richard Brimacombe, Wolfgang Wintermeyer, and Marin van Heel. 1997. “Visualization of Elongation Factor Tu on the Escherichia Coli Ribosome.” Nature 389 (6649): 403–6. 10.1038/38770.

Steffen, Kristan K., Vivian L. MacKay, Emily O. Kerr, et al. 2008a. “Yeast Life Span Extension by Depletion of 60s Ribosomal Subunits Is Mediated by Gcn4.” Cell 133 (2): 292–302. 10.1016/j.cell.2008.02.037.

Uchiumi, T., A. J. Wahba, and R. R. Traut. 1987. “Topography and Stoichiometry of Acidic Proteins in Large Ribosomal Subunits from Artemia Salina as Determined by Crosslinking.” Proceedings of the National Academy of Sciences of the United States of America 84 (16): 5580–84. 10.1073/pnas.84.16.5580.

Wei, Min, Paola Fabrizio, Jia Hu, et al. 2008. “Life Span Extension by Calorie Restriction Depends on Rim15 and Transcription Factors Downstream of Ras/PKA, Tor, and Sch9.” PLoS Genetics 4 (1): e13. 10.1371/journal.pgen.0040013.

Wu, Ziyun, Shao Quan Liu, and Dejian Huang. 2013. “Dietary Restriction Depends on Nutrient Composition to Extend Chronological Lifespan in Budding Yeast Saccharomyces Cerevisiae.” PLoS ONE 8 (5): e64448. 10.1371/journal.pone.0064448.

Yerlikaya, Seda, Madeleine Meusburger, Romika Kumari, et al. 2016. “TORC1 and TORC2 Work Together to Regulate Ribosomal Protein S6 Phosphorylation in Saccharomyces Cerevisiae.” Molecular Biology of the Cell 27 (2): 397–409. 10.1091/mbc.E15-08-0594.

